# The aryl hydrocarbon receptor in β-cells mediates the effects of TCDD on glucose homeostasis in mice

**DOI:** 10.1101/2023.08.10.552760

**Authors:** Myriam P Hoyeck, Ma. Enrica Angela Ching, Lahari Basu, Kyle Van Allen, Jana Palaniyandi, Ineli Perera, Emilia Poleo-Giorgani, Antonio A Hanson, Jennifer E Bruin

**Affiliations:** Department of Biology & Institute of Biochemistry, Carleton University, Ottawa, ON, Canada

**Keywords:** Aryl hydrocarbon receptor, β-cells, cre-loxP knockout, type 2 diabetes, dioxins, environmental pollutants

## Abstract

Chronic exposure to persistent organic pollutants (POPs) is associated with increased incidence of type 2 diabetes, hyperglycemia, and poor insulin secretion in humans. Dioxins and dioxin-like compounds are a broad class of POPs that exert cellular toxicity through activation of the aryl hydrocarbon receptor (AhR). We previously showed that a single high-dose injection of 2,3,7,8-tetrachlorodibenzo-*p*-dioxin (TCDD, aka dioxin; 20 µg/kg) *in vivo* reduced fasting and glucose-stimulated plasma insulin levels for up to 6 weeks in male and female mice. TCDD-exposed male mice were also modestly hypoglycemic and had increased insulin sensitivity, whereas TCDD-exposed females were transiently glucose intolerant; whether these effects are driven by AhR activation in β-cells requires investigation. Here we exposed female and male β-cell specific AhR knockout (βAhr^KO^) mice and littermate Ins1-Cre genotype controls (βAhr^WT^) to a single high dose of 20 µg/kg TCDD and tracked the mice for 6 weeks. We found that deleting AhR from β-cells increased insulin secretion *ex vivo* in female mouse islets and promoted modest weight gain in male mice under baseline conditions. Importantly, high-dose TCDD exposure impaired glucose homeostasis and β-cell function in βAhr^WT^ mice, but these phenotypes were largely abolished in TCDD-exposed βAhr^KO^ mice. Our study demonstrates that AhR signaling in β-cell is important for regulating baseline β-cell function in female mice and energy homeostasis in male mice. We also show that β-cell AhR signaling largely mediates the effects of TCDD on glucose homeostasis in both female and male mice, suggesting that the effects of TCDD on β-cell function/health are driving metabolic phenotypes in peripheral tissues.

## 1. Introduction

Persistent organic pollutants (POPs) are a diverse family of lipophilic environmental contaminants that resist degradation, leading to their widespread global dispersion. Chronic exposure to POPs is associated with numerous adverse health outcomes in humans, including increased risk of type 2 diabetes (T2D) [1–7], hyperglycemia [1,7–11] and poor insulin secretion [1,10–14]. However, the causal link between POP exposure and metabolic dysregulation remains poorly understood.

Dioxins and dioxin-like compounds are a broad class of POPs that exert cellular toxicity through activation of the aryl hydrocarbon receptor (AhR). AhR is a ligand-dependent nuclear receptor that is involved in numerous biological processes. The canonical AhR pathway is involved in the detoxification of xenobiotic compounds (such as dioxins) via activation of phase I (e.g. cytochrome P450, CYP450) and phase II (e.g. NAD(P)H: quinone oxidoreductase, NQO1; glutathione S-transferase alpha 1, GSTA1) xenobiotic metabolism enzymes [15,16]; however, activation of these enzymes can also lead to oxidative stress and consequently DNA damage. Global and tissue-specific AhR knockout models have revealed an essential role of AhR in non-canonical processes, including cell proliferation and adhesion, apoptosis, protein degradation, organ development, immune responses, and circadian rhythms [16–20]. Importantly, global AhR knockout mice have increased insulin sensitivity and improved glucose tolerance on both chow [21] and high-fat diets (HFD) [22], and reduced adiposity when fed a HFD [22], implicating AhR in regulating energy metabolism and glucose homeostasis.

Our lab has shown that direct *in vitro* and systemic *in vivo* exposure to the AhR ligand, 2,3,7,8-tetrachlorodiobenzo-*p*-dioxin (TCDD, aka “dioxin”), induced *Cyp1a1* gene expression in mouse pancreatic islets [23–25]. This indicates that dioxins directly activate AhR within the endocrine pancreas, but the impact of AhR signaling on β-cell function remains unclear. We have shown that a single high-dose injection of TCDD (20 µg/kg) *in vivo* reduced fasting and glucose-stimulated plasma insulin levels for up to 6 weeks in male and female mice [24]. TCDD-exposed male mice were also modestly hypoglycemic and had increased insulin sensitivity, whereas TCDD-exposed females were transiently glucose intolerant compared to CO-exposed mice. It is unclear if the direct action of TCDD on β-cells contributes to these sex-specific metabolic outcomes. Interestingly, high-dose TCDD exposure did not reduced plasma insulin levels in male mice with a global AhR deletion [26], supporting a role of AhR in mediating the effects of TCDD *in vivo*; whether this is driven by the AhR pathway in β-cells requires investigation.

To elucidate the role of AhR in β-cells, we generated a β-cell specific Ahr knockout (βAhr^KO^) mouse model using the Cre-loxP system. We first assessed the role of AhR in regulating body weight, glucose homeostasis, and β-cell health in the absence of exogenous chemical stimuli. We then administered a single high-dose of TCDD (20 µg/kg), as in our previous study [24], to determine whether the effects of TCDD on glucose homeostasis and β-cell function/health in mice are mediated by β-cell AhR signaling. We report that deleting Ahr from β-cells promoted modest weight gain in male mice and increased insulin secretion *ex vivo* in female mouse islets under baseline conditions. Importantly, high-dose TCDD exposure impaired glucose homeostasis and β-cell function in βAhr^WT^ mice (Ins1-Cre genotype control), but these phenotypes were largely abolished in TCDD-exposed βAhr^KO^ mice. Collectively, our study demonstrated that AhR signaling in β-cells largely mediates the effects of TCDD on glucose homeostasis in both female and male mice, most likely by promoting β-cell inflammation and dysfunction.

## 2. Materials and Methods

### 2.1. Animals

All mice received *ad libitum* access to standard rodent chow (Harlan Laboratories, Teklad Diet #2018; Madison, WI) and were maintained on a 12-hour light/dark cycle throughout the study. All experiments were approved by the Carleton University Animal Care Committee and carried out in accordance with the Canadian Council on Animal Care guidelines. Prior to beginning experimental protocols, animals were randomly assigned to treatment groups and matched for body weight and blood glucose to ensure that these variables were consistent between groups.

#### 2.1.1. Generating β-cell specific Ahr knockout mice

B6(Cg)-Ins1^tm1.1(cre)Thor^IJ (#026801; mouse expressing Cre recombinase under the control of the *insulin-1* (*Ins-1*) promoter; **Figure 1C**) and Ahr^tm3.1Bra^IJ (#006203; mouse with loxP sites flanking exon 2 of the *Ahr* allele; **Figure 1B**) were purchased from The Jackson Labs (Bar Harbour, ME, USA). These mice were crossed to generate mice with a partial Ahr knockout in β-cells (βAhr^fl/wt^ Ins1^Cre/wt^), which were further crossed to generate βAhr^fl/fl^ Ins1^wt/wt^ and βAhr^fl/wt^ Ins1^Cre/Cre^ mice (**Figure 1A**). Our experimental mice were then generating by crossing βAhr^fl/fl^ Ins1^wt/wt^ females with βAhr^fl/wt^ Ins1^Cre/Cre^ males (**Figure 1A**). We refer to β-cell specific Ahr knockout mice (βAhr^fl/fl^ Ins1^Cre/wt^) as “βAhr^KO^” and genotype control mice with a single copy of Ins1-Cre (βAhr^wt/wt^ Ins1^Cre/wt^) as “βAhr^WT^” (**Figure 1A**). Genotypes were confirmed by qPCR from an ear notch using primer sequences provided by The Jackson Labs and subsequently running qPCR products on an agarose gel (**Figure 1B,D**; primer sequences in **Supplemental Table 1**).

**Figure 1:**
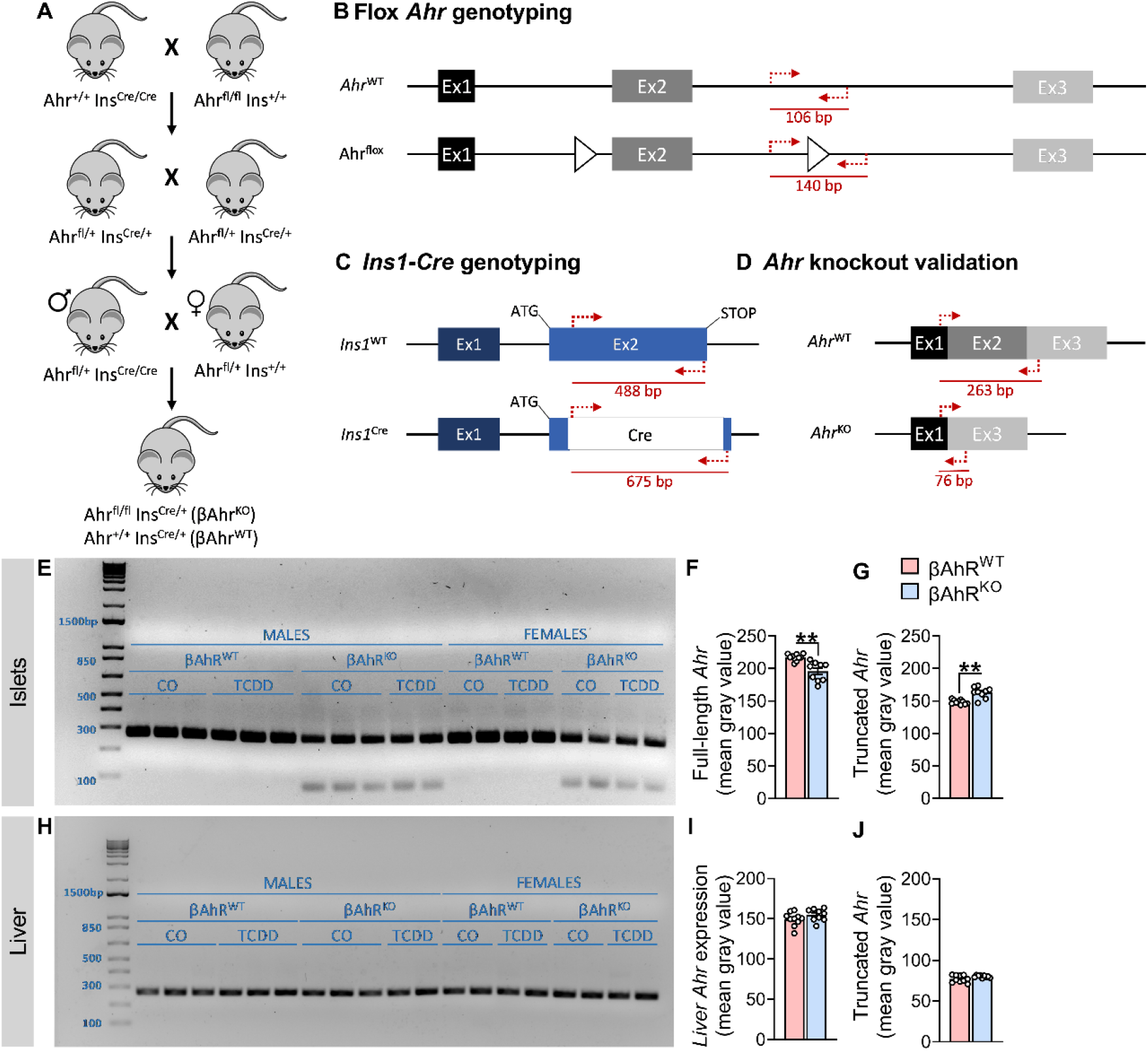
Validation of βAhrKO mouse model. **(A)** Schematic summary of the breeding strategy used to generate experimental mice. Ahr^+/+^ Ins^Cre/Cre^ and Ahr^fl/fl^ Ins^+/+^ mice were crossed to generate heterozygous Ahr^fl/+^ Ins^Cre/+^ mice, which were then crossed to each other to generate Ahr^fl/+^ Ins^Cre/Cre^ and Ahr^fl/+^ Ins^+/+^ mice, which were further crossed to generate Ahr^+/+^ Ins^Cre/+^ (βAhr^WT^) and Ahr^fl/fl^ Ins^Cre/+^ (βAhr^KO^) mice. **(B)** Schematic summary displaying the location of flox sites on the Ahr gene, the binding location of the flox Ahr genotyping primers, and the genotyping product size for Ahr^WT^ and Ahr^flox^ mice. b.p. = base pair. **(C)** Schematic summary of the *insulin-1* gene without Cre (Ins1^WT^) and with Cre (Ins^Cre^), the binding location of the Ins1-Cre genotyping primers, and the genotyping product sizes. **(D)** Schematic summary of the Ahr mRNA in Ahr^WT^ and Ahr^flox^ mice, the binding location of the flox Ahr validation qPCR primers, and the qPCR product size. **(E,H)** Agarose gel of qPCR products showing the presence or absence of *Ahr* recombination in CO and TCDD-exposed βAhr^WT^ and βAhr^KO^ mouse **(E)** islets and **(H)** liver. **(F,G,I,J)** Densitometry analysis of **(F,I)** the full-length *Ahr* product and **(G,J)** the truncated *Ahr* product in βAhr^WT^ and βAhr^KO^ mice seen on the agarose gels in **(E,H)**; sex and chemical treatment were not consider in this analysis and all data was pooled by genotype. **p ≤ 0.01. The following statistical tests were used: **(F,G,J)** two-tailed Mann-Whitney test; **(I)** two-tailed unpaired t-test.

We confirmed deletion of Ahr by verifying that recombination was occurring along exon 2 of the *Ahr* gene in whole islets. Islets were isolated from βAhr^WT^ and βAhr^KO^ mice at 33 – 41 weeks of age, as described in section “***2.3. Islet isolations***”. *Ahr* was amplified using qPCR (**Figure 1D**; primers sequences in **Supplemental Table 2**), as described in “***2.6. Quantitative real time PCR***”, and gene products were run on a 2% agarose gel. The same method was used to ensure that no *Ahr* recombination was occurring in liver of βAhr^KO^ mice.

#### 2.1.2. Experimental protocol

Female and male βAhr^KO^ and βAhr^WT^ mice were generated as described in section “***2.1.1. Generating β-cells specific Ahr knockout mice***”; 45 litters were generated, with ∼58% of litters having both βAhr^KO^ mice and corresponding βAhr^WT^ littermate controls. At 6 – 37 weeks of age, βAhr^KO^ and βAhr^WT^ mice received a single intraperitoneal (i.p.) injection of corn oil (CO; 25 ml/kg, vehicle control, Sigma-Aldrich, #C8267-2.5L; St Louise, MO, USA) or 20 µg/kg TCDD (AccuStandard, #AD404SDMSO10X), as outlined in **Figure 3A** (n = 11 – 13 mice / group). At 1-week post-injection, islets were isolated from a subset of mice for an *ex vivo* glucose-stimulated insulin secretion (GSIS) assay, cell viability assessment, or RNA extraction (n = 6 – 7 mice / group); liver was extracted from the same subset of mice and flash frozen for qPCR analysis (n = 6 – 7 mice / group). Whole pancreas also harvested from another subset of mice at 1-week post-injection and stored in 4% paraformaldehyde (PFA) for 24 hours, followed by long-term storage in 70% ethanol for histological analysis (n = 5 – 7 mice / group). At 6 weeks post-injection, islets were isolated from a third subset of mice for an *ex vivo* GSIS assay and RNA extraction.

### 2.2. Metabolic assessments

All metabolic analyses were performed in conscious, restrained mice, and blood samples were collected via the saphenous vein using heparinized microhematocrit tubes (Fisherbrand, #22-362-566). Blood glucose levels were measured using a handheld glucometer (MediSure; Ajax, Canada). Body weight and blood glucose were measured following a 4-hour morning fast 1-2x/week throughout the study. For all metabolic tests, time 0 indicates the blood sample collected prior to administration of glucose or insulin. For glucose tolerance tests (GTTs), mice received an i.p. bolus of glucose (2 g/kg) following a 6-hour morning fast. Blood samples were collected at 0, 15, 30, and 60 minutes for measuring plasma insulin levels by ELISA (ALPCO, #80-INSMSU-E01; Salem, NH, USA). For insulin tolerance tests (ITTs), mice received an i.p. bolus of insulin (0.7 IU/kg; Novolin ge Toronto, #02024233; Novo Nordisk, Canada) following a 4-hour morning fast.

### 2.3. Islet isolation

Islets were isolated by pancreatic duct injection with collagenase (1,000 units/ml; Sigma Aldrich, #C7657) dissolved in Hanks’ balanced salt solution (HBSS: 137 mM NaCl, 5.4 mM KCl, 4.2 mM NaH_2_PO_4_, 4.1 mM KH_2_PO_4_, 10 mM HEPES, 1 mM MgCl_2_, 5 mM dextrose, pH 7.2). Pancreata were incubated at 37°C for 10 minutes 45 seconds, vigorously agitated, and the collagenase reaction quenched by adding cold HBSS with 1mM CaCl_2_. The pancreas tissue was washed 3 times in HBSS+CaCl_2_ (centrifuging for 1 minute at 1000 rpm between washes) and resuspended in RPMI 1640 1X (Wisent Bioproducts, 350-000-CL; St-Bruno, Canada) containing 10% fetal bovine serum (FBS; Sigma-Aldrich, #F1051-500ml) and 1% 10000 IU/ml penicillin-streptomycin (Gibco, #15140-122-100). Pancreas tissue was filtered through a 70 μm cell strainer and islets were handpicked under a dissecting scope to >95% purity. The day following isolation, islets were used for *ex vivo* GSIS, cell viability assessment or stored in buffer RLT with 1% β-mercaptoethanol for RNA extraction.

### 2.4. Ex vivo glucose-stimulated insulin secretion assay

To assess β-cell function, 25 islets per replicate (n = 3 technical replicates per mouse; n= 3 – 8 mice / group) were transferred to a 1.5 mL microcentrifuge tube and washed with pre-warmed (37°C) Krebs–Ringer bicarbonate buffer (KRBB) with 0 mM glucose (115 mM NaCl, 5 mM KCl, 24 mM NaHCO_3_, 2.5 mM CaCl_2_, 1 mM MgCl_2_, 10 mM HEPES, 0.1% (wt/vol.) BSA, pH 7.4). Islets were then immersed in KRBB with 2.8 mM glucose (low glucose, LG) for a 1-hour pre-incubation at 37°C and the supernatant was discarded. Islets were then immersed in 500 μl of LG KRBB for 1 hour, followed by 500 μl of KRBB with 16.7 mM glucose (high glucose, HG) for 1 hour at 37°C. The LG KRBB and HG KRBB samples were centrifuged (2000 rpm) and the supernatant stored at −30°C until use. To measure insulin content, islets were immersed in 500 µl of acid-ethanol solution (1.5% (vol./vol.) HCl in 70% (vol./vol.) ethanol) at 4°C overnight and then neutralised with 1 M Tris base (pH 7.5) before long-term storage at −30°C. Insulin concentrations were measured by ELISA (ALPCO, #80-INSMR).

### 2.5. Islet cell viability

To assess cell viability, 50 islets per replicate (n = 2 technical replicates per mouse; n = 3 – 9 mice / group) were transferred to a 1.5 mL microcentrifuge tube and washed with pre-warmed PBS+/+ (Sigma, # D8662). Islets were dispersed by adding 400 µl Accutase (VWR, CA10215-416) and incubating islets at 37°C for ∼4 minutes with trituration occurring every 2 minutes (VWR, #CA10215-416). Accutase was neutralized with 600 µl RPMI 1640 + 10% FBS + 1% 10000 IU/ml penicillin-streptomycin. Cells were washed once with PBS+/+ and then stained with 0.5 µM Hoescht (Thermo Scientific, #62249), 1.25 µM Calcein (Invitrogen, #L3224), and 0.75 µM PI (Invitrogen, #P21493) at room temperature for 30 minutes in a 96-well flat-bottom plate pre-coated with poly-D-lysine (Sigma-Aldrich #P7280). An Axio Observer 7 microscope was used to image 10% of each well immediately after staining. The number of calcein^+^ cells (live) and PI^+^ cells (dead/dying) was quantified using Zen Blue 2.6 software (Carl Zeiss, Germany). The % live cell was calculated as [(# of live cells imaged in well) / (total # cell images in well) *100], with an average of 1650 cells counted per well.

### 2.6. Quantitative real time PCR

RNA was extracted from isolated islets stored in buffer RLT with 1% β-mercaptoethanol using the Qiagen RNeasy Micro Kit (#74004), as per the manufacturer’s instructions. RNA was extracted from flash frozen liver using TRIzol^TM^ (Invitrogen, #15596018; Carlsbad, CA, USA), as per the manufacturer’s instructions. DNase treatment was performed prior to cDNA synthesis with the iScript^TM^ gDNA Clear cDNA Synthesis Kit (Biorad, #1725035; Mississauga, ON, Canada). qPCR was performed using SsoAdvanced Universal SYBR Green Supermix (Biorad, #1725271) and run on a CFX384 (Biorad). *PPIA* was used as the reference gene since this gene displayed stable expression under control and treatment conditions. Data were analyzed using the 2^-ΔΔCT^ relative quantitation method. Primer sequences are listed in **Supplemental Table 1 – 3**.

### 2.7. Immunofluorescence staining and image quantification

PFA-fixed pancreas tissues were processed and paraffin-embedded by the University of Ottawa Heart Institute Histology Core Facility (Ottawa, ON). Immunofluorescent staining was performed on tissue sections (5 µm thick), as previously described [23]. Apoptosis was assessed using the Molecular Probes Click-It Plus TUNEL Assay with Alexa Fluor 488 dye (Invitrogen, #C10617), according to manufacturer’s instructions. Pancreas sections were counterstained with rabbit anti-insulin (1:200, C27C9; Cell Signaling Technology, #3014, Danvers, MA, USA) and goat anti-rabbit IgG (H+L) secondary antibody, Alexa Fluor 594 (1:1000; Invitrogen, #A11037) with no antigen retrieval. For every round of staining, a pancreas section was treated with DNase (Qiagen, # 1023460) prior to TUNEL staining as a positive control.

For islet morphology quantification, the entire pancreas section was imaged with an Axio Observer 7 microscope and all islets within the field of view were quantified. Immunofluorescence was manually quantified using Zen Blue 2.6 software. The average of all islet measurements is reported for each biological replicate. The % INS^+^ area per islet was calculated as [(hormone^+^ area / islet area) x 100], with a range of 2 – 8 islets quantified per mouse. The % TUNEL^+^ INS^+^ cells per pancreas section was calculated as [(# of TUNEL^+^ INS^+^ cells per pancreas section) / (total # of INS^+^ cells per pancreas section) x 100], with an average of 350 cells counted per pancreas section.

### 2.8. Quantification and statistical analysis

All statistics were performed using GraphPad Prism 9.5.1 (GraphPad Software Inc., La Jolla, CA). Specific statistical tests are indicated in figure legends. Sample sizes are described in section “***2.1 Animals***” and in figure legends. For all analyses, p ≤ 0.05 was considered statistically significant. Statistically significant outliers were detected by a Grubbs’ test with α = 0.05. All data was tested for normality using a Shapiro-Wilk test and for equal variance using either a Brown-Forsyth test (for one-way ANOVAs) or an F test (for unpaired t-tests). Non-parametric statistics were used in cases where the data failed normality or equal variance tests. Parametric tests were used for all two-way ANOVAs, but normality and equal variance were tested on area under the curve values and by one-way ANOVAs. Data in line and bar graphs display mean ± SEM. Individual data points on bar plots are always biological replicates (i.e. different mice).

## 3. Results

### 3.1. Validation and characterization of βAhr^KO^ mice

Our breeding scheme involved crossing Ahr^+/+^ Ins^Cre/Cre^ and Ahr^fl/fl^ Ins^+/+^ mice to generate heterozygous Ahr^fl/+^ Ins^Cre/+^ mice, which were then crossed to each other to generate Ahr^fl/+^ Ins^Cre/Cre^ and Ahr^fl/+^ Ins^+/+^ mice (**Figure 1A**). These mice were further crossed to generate Ahr^+/+^ Ins^Cre/+^ (“βAhr^WT^”) and Ahr^fl/fl^ Ins^Cre/+^ (“βAhr^KO^”) littermates (**Figure 1A**). We first validated our βAhr^KO^ model by assessing *Ahr* recombination in whole islets by qPCR analysis (**Figure 1D**). A single *Ahr* product band (263 base pairs, bp) was observed in βAhr^WT^ islets (**Figure 1E**) and liver (**Figure 1H**), indicating the presence of exon 2. CO and TCDD-exposed βAhr^KO^ mice had a truncated *Ahr* gene product (76 bp) (**Figure 1E,H**) and a decrease in expression of the full-length *Ahr* gene in islets (**Figure 1F**) but not liver (**Figure 1I-J**), confirming that *Ahr* recombination is specific to islets. The presence of a full-length *Ahr* gene product in βAhr^KO^ islets (**Figure 1E**) was expected given that islets are a mixture of β-cells and other endocrine cells.

We first assessed whether Ahr deletion in β-cells impacted metabolic health under baseline conditions (i.e. in the absence of exogenous chemical exposure). βAhr^KO^ females had normal body weight (**Figure 2A**) and normal fasted plasma insulin levels (**Figure 2C**), but a modest decrease in fasted blood glucose levels (**Figure 2B – AUC**) compared to βAhr^WT^ females. There was no effect of genotype on overall fasted blood glucose (**Figure 2E – AUC**) or plasma insulin levels (**Figure 2F**) in male mice. Interestingly, βAhr^KO^ males had significantly higher body weight compared to βAhr^WT^ males (**Figure 2D**). These data suggest that AhR signaling in β-cells plays a role in maintaining metabolic health under baseline conditions.

**Figure 2:**
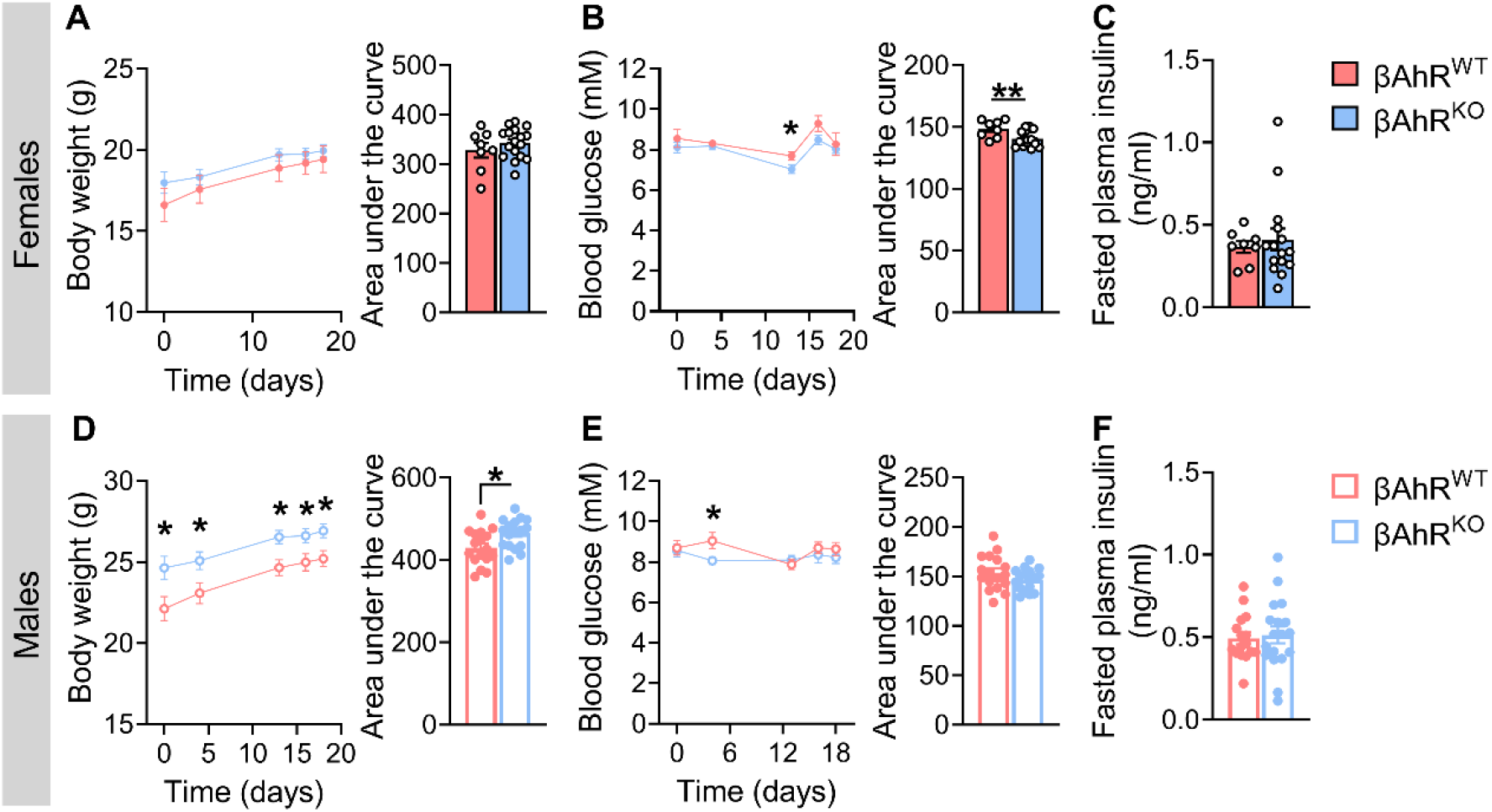
Deleting Ahr in β-cells causes hypoglycemia in females and increased body weight in males under baseline condition. Mice were 4 – 8 weeks old when tracking was started at t = 0 days. **(A,D)** Body weight, **(B,E)** fasted blood glucose, and **(C,F)** fasted plasma insulin in **(A-C)** female and **(D-F)** male βAhr^WT^ and βAhr^KO^ mice. All data are presented as mean ± SEM. Individual data points on bar graphs represent biological replicates (different mice). *p ≤ 0.05. The following statistical tests were used: **(A)** line graph, two-way REML ANOVA with uncorrected Fisher’s LSD test; bar graphs, two-tailed unpaired t-test; **(B,D,E)** line graph, two-way REML ANOVA with uncorrected Fisher’s LSD test; bar graphs, Mann-Whitney test; **(C,F)** two-tailed unpaired t-test.

### 3.2. Deleting Ahr from β-cells protected TCDD-exposed male mice from fasting hypoinsulinemia and TCDD-exposed female mice from transient hypoglycemia

We next assessed whether AhR in β-cells mediates the effects of TCDD on glucose homeostasis and β-cell function. Female and male βAhr^WT^ and βAhr^KO^ mice were exposed to CO or a single high dose of TCDD (20 µg/kg) and tracked for up to 6 weeks post CO/TCDD injection (**Figure 3A**). TCDD exposure had no effect on body weight in female mice (**Figure 3B**), or body weight (**Figure 3C**) and fasting blood glucose (**Figure 3G,H**) in male mice, irrespective of genotype. Interestingly, TCDD caused transient hypoglycemia 1-day post-injection in βAhr^WT^ but not βAhr^KO^ females (**Figure 3D,E**).

**Figure 3:**
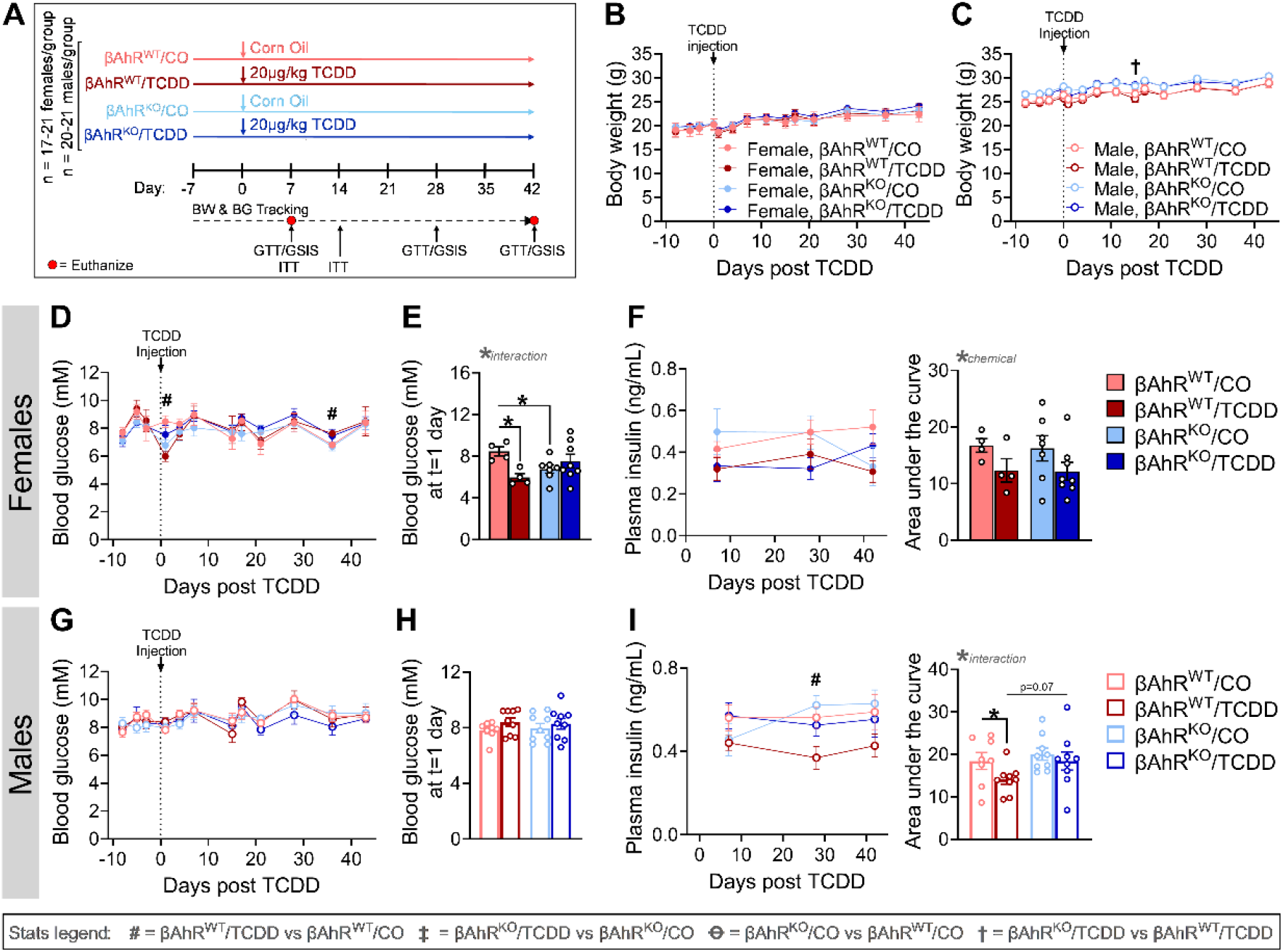
Deleting Ahr from β-cells protected TCDD-exposed male mice from fasting hypoinsulinemia and TCDD-exposed female mice from transient hypoglycemia. **(A)** Schematic summary timeline of the study. Female and male mice were injected with either corn oil (CO) or a single high dose of TCDD (20 µg/kg) and tracked for up to 6 weeks post TCDD/CO injection. A subset of mice was euthanized at week 1 week (n = 11 – 13 / group) and week 6 (n = 4 – 9 / group). BW = body weight, BG = blood glucose, GSIS = glucose-stimulated insulin secretion, GTT = glucose tolerance test, ITT = insulin tolerance test, βAhr^WT^ = Ins^Cre/+^Ahr^+/+^, βAhr^KO^ = Ins^Cre/+^Ahr^fl/fl^. **(B,C)** Body weight, **(D,G)** fasted blood glucose, and **(F,I)** fasted plasma insulin levels were measured weekly in **(B,D,F)** female and **(C,G,I)** male mice. **(E,H)** fasted blood glucose at 1-day post-injection in **(E)** female and **(H)** male mice. All data are presented as mean ± SEM. Individual data points on bar graphs represent biological replicates (different mice). Stats legend (p ≤ 0.05): #, βAhr^WT^/TCDD vs βAhr^WT^/CO; ‡, βAhr^KO^/TCDD vs βAhr^KO^/CO; Ꝋ, βAhr^KO^/CO vs βAhr^WT^/CO; †, βAhr^KO^/TCDD vs βAhr^KO^/TCDD. The following statistical tests were used: **(B,C,D,G)** two-way RM ANOVA with uncorrected Fisher’s LSD test; **(E,H)** two-way ANOVA with uncorrected Fisher’s LSD test; **(F,I)** line graph, two-way RM ANOVA with uncorrected Fisher’s LSD test; bar graphs, two-way ANOVA with Tukey’s multiple comparison test.

We were particularly interested in whether βAhr^KO^ mice would be protected from the TCDD-induced hypoinsulinemia observed in WT mice from our previous study [24]. TCDD caused an overall decrease in fasted plasma insulin levels in females throughout our 6-week study, irrespective of genotype (**Figure 3F – AUC**). Interestingly, TCDD-exposed βAhr^WT^ males displayed significant hypoinsulinemia at 4-weeks post-injection and a trending decrease at 1- and 6-weeks post-injection compared to CO-exposed βAhr^WT^ males, whereas TCDD-exposed βAhr^KO^ males had normal fasted plasma insulin throughout the study (**Figure 3I**). These data imply that AhR signaling in β-cells is mediating the effects of TCDD on fasted plasma insulin levels in male mice.

### 3.3. Deletion of Ahr in β-cells protected female and male mice from TCDD-induced changes in glycemia

To determine the impact of β-cell AhR on regulating glucose homeostasis following TCDD exposure, we performed ipGTTs at 1-, 4-, and 6-weeks post CO/TCDD injection. TCDD exposure had no overall effect on glucose tolerance in βAhr^WT^ or βAhr^KO^ females at 1-week post-injection (**Figure 4A,B**). However, by 4 weeks post-injection, TCDD-exposed βAhr^WT^ females had modest glucose intolerance (**Figure 4Cii,D**), which became more pronounced by 6 weeks post-injection (**Figure 4Eii,F**). TCDD-exposed βAhr^KO^ females maintained normal glycemia relative to CO-exposed females at both week 4 and 6 (**Figure 4Ciii,D,Eiii, F**).

**Figure 4:**
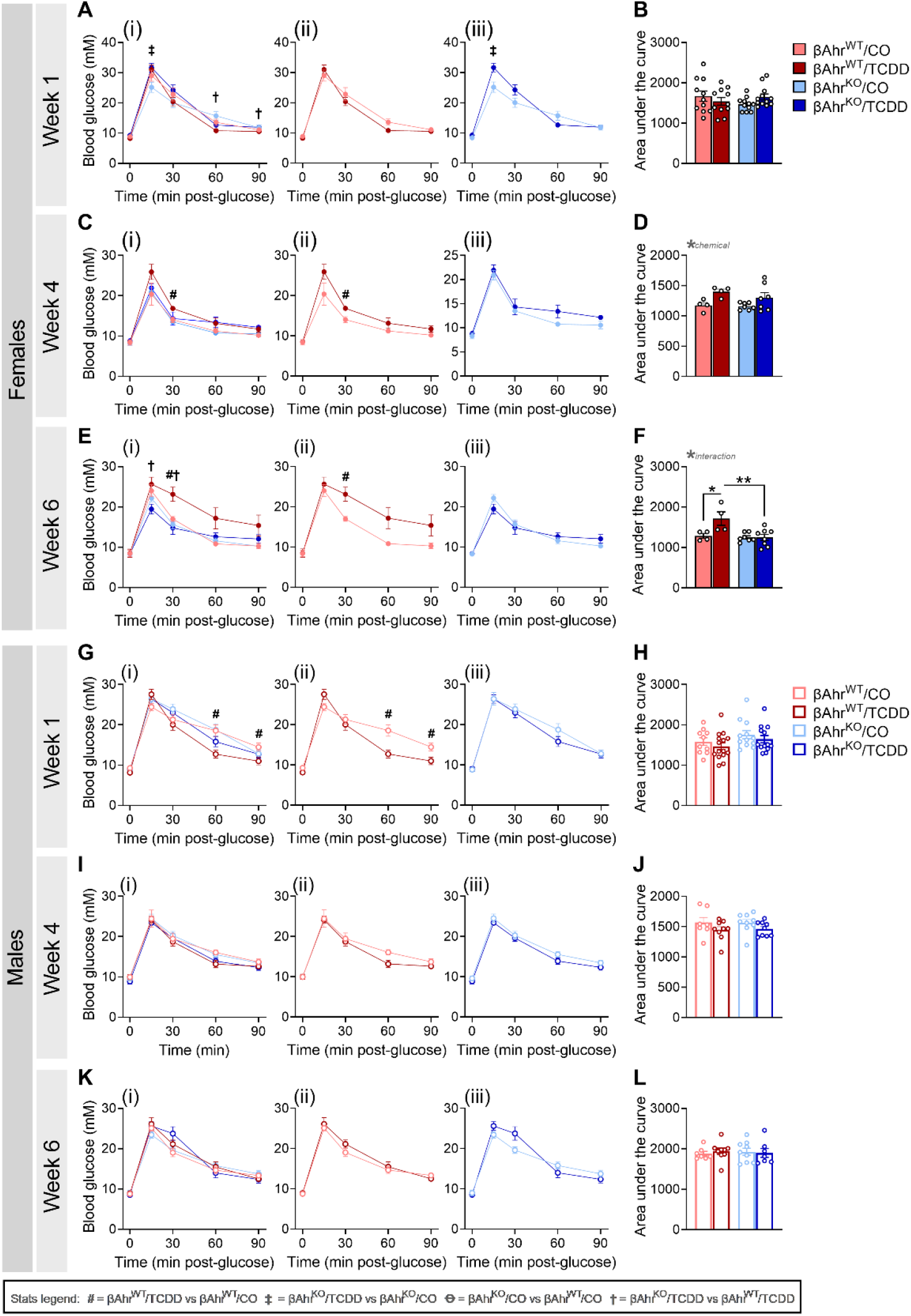
Deletion of Ahr in β-cells protected female and male mice from TCDD-induced changes in glycemia. Glucose tolerance tests were performed at **(A,B,G,H)** 1 week, **(C,D,I,J)** 4 weeks, and **(E,F,K,L)** 6 weeks post TCDD/CO injection (see Figure 3A for study timeline). Data is presented as a line graph and area under the curve. Line graphs are presented as **(i)** all treatment groups, **(ii)** βAhr^WT^/TCDD vs βAhr^WT^/CO, and **(iii)** βAhr^KO^/TCDD vs βAhr^KO^/CO. All data are presented as mean ± SEM. Individual data points in bar graphs represent biological replicates (different mice; n = 4 – 13 mice / group). Stats legend (p ≤ 0.05): #, βAhr^WT^/TCDD vs βAhr^WT^/CO; ‡, βAhr^KO^/TCDD vs βAhr^KO^/CO; Ꝋ, βAhr^KO^/CO vs βAhr^WT^/CO; †, βAhr^KO^/TCDD vs βAhr^KO^/TCDD. The following statistical tests were used: line graph, two-way RM ANOVA with uncorrected Fisher’s LSD test; bar graphs, two-way ANOVA with Tukey’s multiple comparison test.

TCDD-exposed βAhr^WT^ males were significantly hypoglycemic compared to CO-exposed βAhr^WT^ males at 1-week post-injection (**Figure 4Gii**), but normoglycemia was restored by 4 weeks (**Figure 4I–L**). In contrast, TCDD exposure had no effect on glucose tolerance in βAhr^KO^ males (**Figure 4Giii –Liii**). These data imply that AhR signaling in β-cells mediates the effects of TCDD on glucose tolerance.

### 3.4. TCDD exposure caused glucose-stimulated hypoinsulinemia in βAhr^WT^ but not βAhr^KO^ female and male mice

Consistent with the GTT data, TCDD exposure had no overall effect on plasma insulin levels in females of either genotype at 1 week (**Figure 5A,B**) or 4 weeks (**Figure 5C,D**) post-injection. However, by 6 weeks, TCDD-exposed βAhr^WT^ females displayed a trending decrease in plasma insulin levels (**Figure 5Eii,F**), which was not observed in βAhr^KO^ females (**Figure 5Eiii,F**). TCDD exposure caused severe hypoinsulinemia in βAhr^WT^ males compared to CO-exposed βAhr^WT^ males at 1-week post-exposure (**Figure 5Gii,H**), with the severity of hypoinsulinemia diminishing over time (**Figure 5Iii,J,Kii,L**); this phenotype was abolished by knocking out Ahr in β-cells (**Figure 5G–L).**

**Figure 5:**
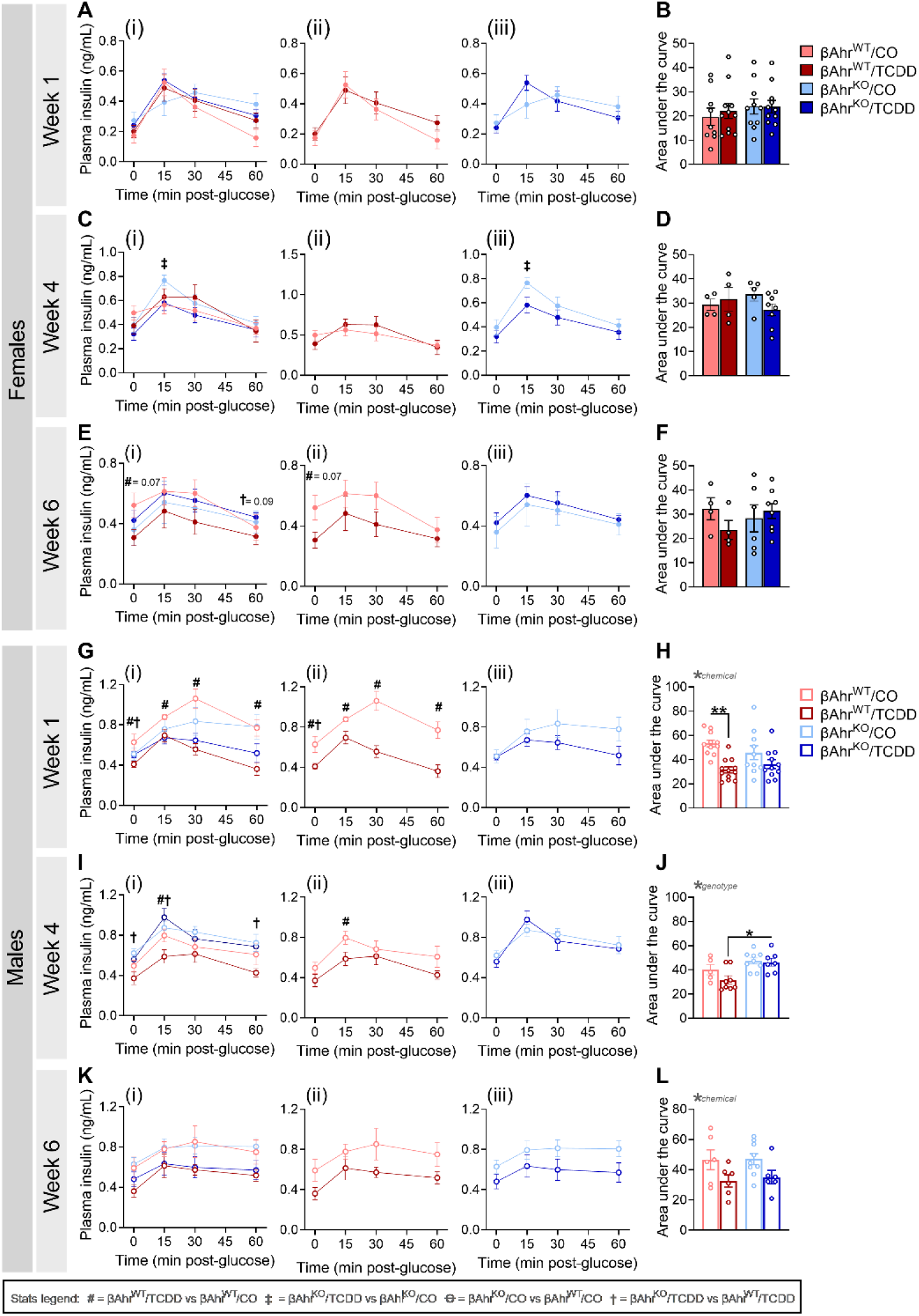
TCDD exposure caused glucose-stimulated hypoinsulinemia in βAhrWT but not βAhrKO female and male mice. Plasma insulin levels were measured during glucose tolerance tests at **(A,B,G,H)** 1 week, **(C,D,I,J)** 4 weeks, and **(E,F,K,L)** 6 weeks post TCDD/CO injection (see Figure 3A for study timeline). Data are presented as a line graph and area under the curve. Line graphs are presented as **(i)** all treatment groups, **(ii)** βAhr^WT^/TCDD vs βAhr^WT^/CO, and **(iii)** βAhr^KO^/TCDD vs βAhr^KO^/CO. All data are presented as mean ± SEM. Individual data points in bar graphs represent biological replicates (different mice; n= 4 – 12 mice / group). Stats legend (p ≤ 0.05): #, βAhr^WT^/TCDD vs βAhr^WT^/CO; ‡, βAhr^KO^/TCDD vs βAhr^KO^/CO; Ꝋ, βAhr^KO^/CO vs βAhr^WT^/CO; †, βAhr^KO^/TCDD vs βAhr^KO^/TCDD. The following statistical tests were used: line graph, two-way RM ANOVA with uncorrected Fisher’s LSD test; bar graphs, two-way ANOVA with Tukey’s multiple comparison test.

In summary, our GTT data shows that Ahr deletion in β-cells had little to no effect on glucose homeostasis or glucose-stimulated plasma insulin levels in female and male mice exposed to corn oil. However, β-cell AhR plays a clear role in the metabolic response to an exogenous AhR ligand.

### 3.5. βAhr^KO^ female and male mice were protected from TCDD-induced changes in insulin sensitivity

We previously reported that TCDD-exposed male WT mice were more insulin sensitive than CO-exposed males [24]. Since changes in insulin secretion can drive adaptations in peripheral sensitivity [27,28], we examined whether β-cell Ahr deletion mitigated the effects of TCDD on insulin sensitivity (**Figure 6A,B,E,F**). In βAhr^WT^ females, TCDD exposure caused a modest increase in insulin sensitivity at 15-minutes post-insulin, followed by insulin resistance at 60- and 90-minutes post-insulin compared to CO-injected controls (**Figure 6Aii**); this effect was largely absent in TCDD-exposed βAhr^KO^ females (**Figure 6Aiii**). TCDD-exposed βAhr^WT^ males were significantly more insulin sensitive throughout the ITT compared to CO-exposed βAhr^WT^ males (**Figure 6Eii**), whereas TCDD had no effect on insulin sensitivity in βAhr^KO^ males (**Figure 6Eiii**).

**Figure 6:**
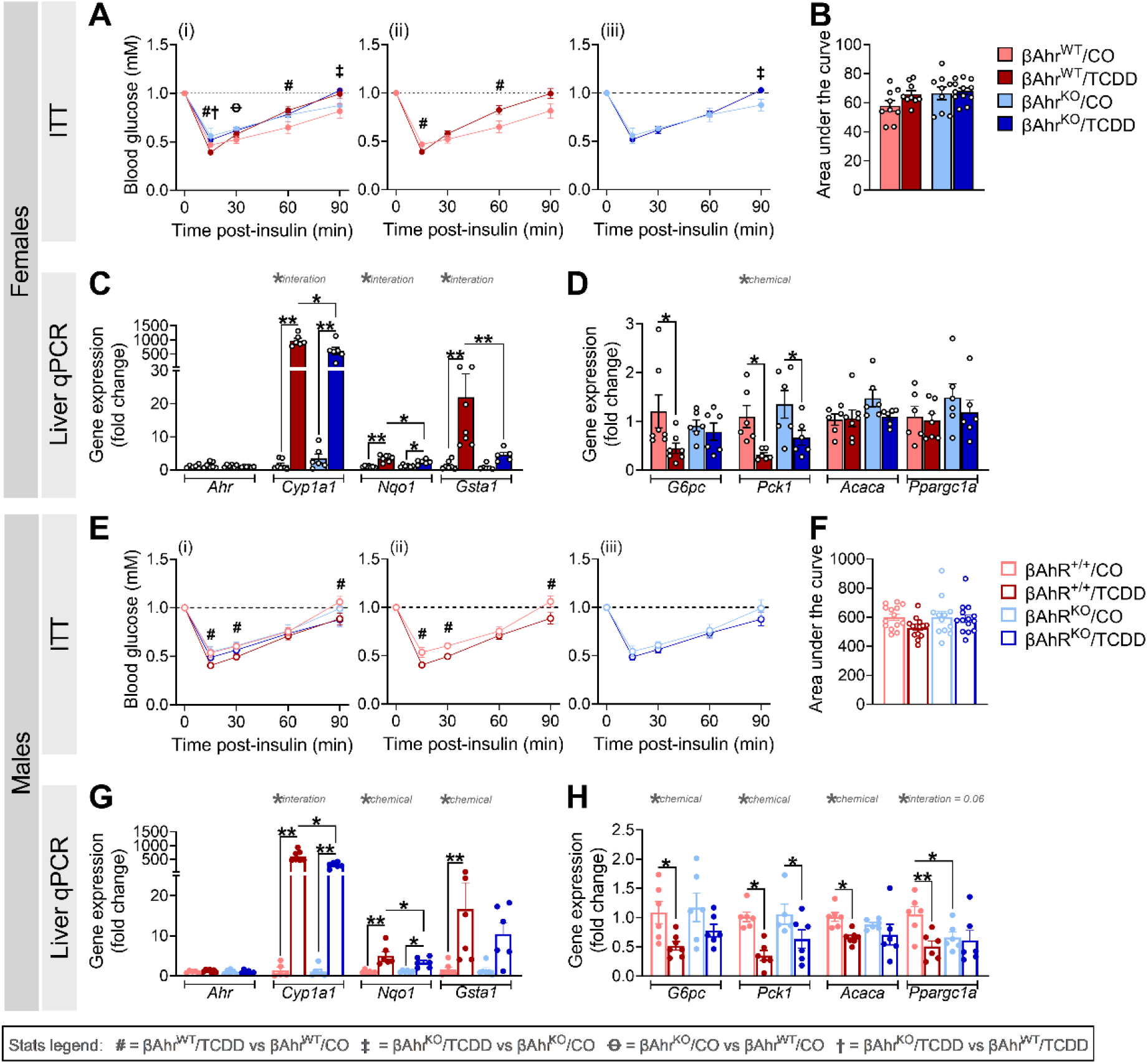
βAhrKO female and male mice were protected from TCDD-induced changes in insulin sensitivity. An insulin tolerance test (ITT) was performed at 1-2 weeks post TCDDD/CO injection, and liver was harvested at 1-week post TCDD/CO injection to assess gene expression by qPCR (see Figure 3A for study timeline). Blood glucose levels during the ITT in **(A,B)** female and **(E,F)** male mice (n = 9 – 14 mice / group). **(C,G)** Markers of the AhR pathway in liver from **(C)** females and **(G)** males. **(D,H)** Insulin-dependent markers of gluconeogenesis and lipogenesis (n = 5 – 7 mice / group). ITT data is presented as a line graph and area under the curve; line graphs are presented as **(i)** all treatment groups, **(ii)** βAhr^WT^/TCDD vs βAhr^WT^/CO, and **(iii)** βAhr^KO^/TCDD vs βAhr^KO^/CO. All data are presented as mean ± SEM. Individual data points in bar graphs represent biological replicates (different mice). *p ≤ 0.05, **p ≤ 0.01. Stats legend (p ≤ 0.05): #, βAhr^WT^/TCDD vs βAhr^WT^/CO; ‡, βAhr^KO^/TCDD vs βAhr^KO^/CO; Ꝋ, βAhr^KO^/CO vs βAhr^WT^/CO; †, βAhr^KO^/TCDD vs βAhr^KO^/TCDD. The following statistical tests were used: **(A,E)** two-way RM ANOVA with uncorrected Fisher’s LSD test; **(B,F)** two-way ANOVA with Tukey’s multiple comparison test; **(C,D,G,H**) two-way ANOVA with uncorrected Fisher’s LSD test.

The ITT results suggest a role for β-cell AhR signaling in driving the effects of TCDD on peripheral tissues. To further explore this idea, we measured gene expression for markers of the AhR pathway and insulin signaling in liver tissue from βAhr^WT^ and βAhr^KO^ mice (**Figure 6C,D,G,H**). There were no changes in liver *Ahr* expression, irrespective of genotype or chemical exposure (**Figure 6C,G**). Interestingly, *Cyp1a1*, *Nqo1*, and *Gsta1* expression were significantly upregulated in liver of TCDD-exposed βAhr^WT^ and βAhr^KO^ mice compared to CO-exposed mice, but expression was dampened in TCDD-exposed βAhr^KO^ liver compared to βAhr^WT^ liver (**Figure 6C,G**) despite the absence of Ahr recombination in the liver (**Figure 1H–J**).

TCDD exposure also reduced expression of the gluconeogenesis enzyme *G6pc* (glucose-6-phosphatase) in liver from βAhr^WT^ female (**Figure 6D**) and male (**Figure 6H**) mice compared to CO exposure, an effect that was absent in βAhr^KO^ mice (**Figure 6D,H**). Similarly, the lipogenesis markers *Acaca* (acetyl-CoA carboxylase alpha; **Figure 6H**) and *Ppargc1a* (peroxisome proliferator-activated receptor gamma coactivator 1-alpha; **Figure 6H**) was reduced in TCDD-exposed βAhr^WT^ males compared to CO-exposed βAhr^WT^ males, but Ahr deletion in β-cells prevented this effect. In contrast, TCDD reduced expression of the gluconeogenesis enzyme *Pck1* (phosphoenolpyruvate carboxykinase 1) in female (**Figure 6D**) and male (**Figure 6H**) liver, irrespective of genotype. There was no effect of chemical or genotype on liver *Acaca* or *Ppargc1a* expression in females (**Figure 6D**). Collectively, these data suggest that the effects of TCDD on β-cell physiology are driving changes in peripheral insulin sensitivity, most likely due to AhR-mediated changes in insulin secretion.

### 3.6. TCDD exposure impaired insulin secretion ex vivo in βAhr^WT^ but not βAhr^KO^ male islets

Next, we assessed whether AhR signaling in β-cells impacts β-cell function by measuring GSIS in isolated islets *ex vivo* at 1- and 6-weeks following CO/TCDD injection. Interestingly, βAhr^KO^ female islets showed increased basal insulin secretion under LG conditions compared to βAhr^WT^ islets (**Figure 7B,F**). βAhr^KO^ female islets also had increased insulin secretion under high-glucose condition compared to βAhr^WT^ islets at week 6 (**Figure 7E**), pointing to a role of AhR in regulating β-cell function in female mice.

**Figure 7:**
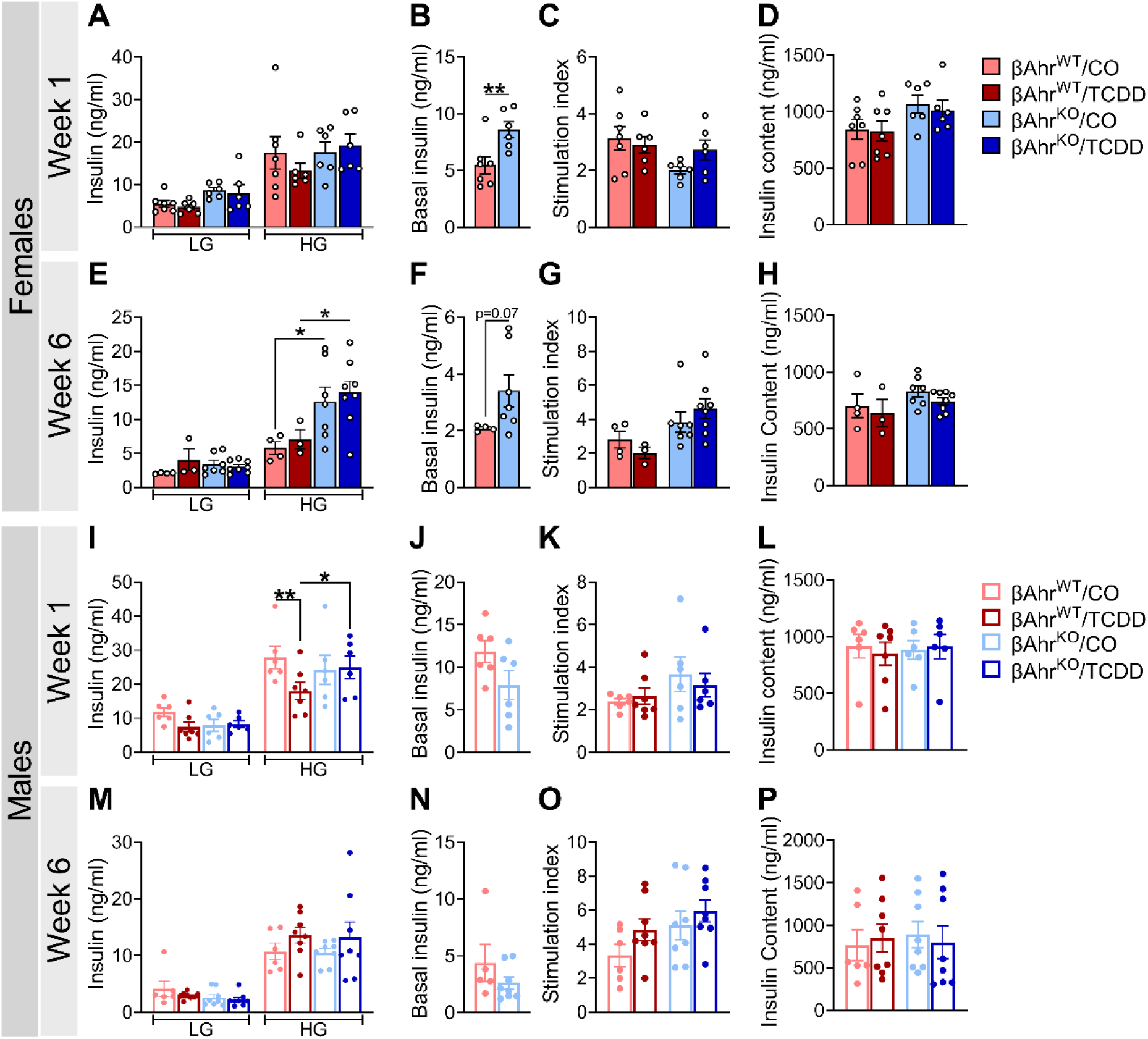
TCDD exposure impaired insulin secretion *ex vivo* in βAhrWT but not βAhrKO male islets. Islets were isolated at 1 and 6 weeks post TCDD/CO injection to assess islet function *ex vivo* (see Figure 3A for study timeline). LG = 2.8 mM low glucose, HG = 16.7 mM high glucose. **(A,E,I,M)** Insulin secretion following a sequential 1-hour incubation in LG and HG buffer at **(A,I)** week 1 and **(E,M)** week 6 in **(A,E)** female and **(I,M)** male mice. **(B,F,J,N)** Basal insulin secretion following a 1-hour incubation in LG buffer in CO-exposed βAhr^WT^ and βAhr^KO^ **(B,F)** female and **(J,N)** male mice at **(B,J)** week 1 and **(F,N)** week 6. **(C,G,K,O)** Stimulation index at **(C,K)** week 1 and **(G,O)** week 6 in **(C,G)** female and **(K,O)** male mice; stimulation index was calculated as a ratio of insulin concentration under HG condition relative to LG. **(D,H,L,P)** Insulin content of islets following an overnight incubation in acid ethanol at **(D,L)** week 1 and **(H,P)** week 6 in **(D,H)** female and **(L,P)** male mice. All data are presented as mean ± SEM. Individual data points in bar graphs represent biological replicates (different mice; n= 3 – 8 mice / group), with each biological replicate representing an average of three technical replicates. *p ≤ 0.05, **p ≤ 0.01. The following statistical tests were used: **(A,C– E,G–I,K–M,O,P)** two-way ANOVA with uncorrected Fisher’s LSD test; **(B,F,J,N)** two-tailed unpaired t-test.

TCDD had no effect on GSIS (**Figure 7A,E**), stimulation index (**Figure 7C,G**), or insulin content (**Figure 7D,H**) in female islets at 1- or 6-weeks post-injection, irrespective of genotype. In contrast, high-dose TCDD exposure *in vivo* significantly impaired *ex vivo* GSIS in βAhr^WT^ male islets but not βAhr^KO^ male islets at 1-week post-injection (**Figure 7I**); GSIS was restored in TCDD-exposed βAhr^WT^ male islets by 6 weeks post-injection (**Figure 7M**). TCDD exposure had no effect on stimulation index (**Figure 7K,O**) or insulin content (**Figure 7L,P**) in male islets, irrespective of genotype. These data suggest that AhR signaling in β-cells is driving TCDD-induced β-cell dysfunction in males and likely explains the pronounced hypoinsulinemia in βAhr^WT^ males *in vivo* following TCDD exposure (**Figure 5G–J**).

Collectively, our data imply that AhR expression in β-cells plays a role in maintaining basal insulin secretion in female islets under non-chemical conditions. TCDD exposure *in vivo* impairs β-cell function in male but not female βAhr^WT^ islets, and knocking out β-cell AhR mitigated this effect.

### 3.7. TCDD exposure increased Il1b expression in βAhr^WT^ but not βAhr^KO^ female islets

We assessed the effects of *in vivo* TCDD exposure on islet cell apoptosis and inflammation in isolated islets at 1-week post-injection. There was no effect of chemical or genotype on *Tnfα* or *Nf-Κβ* expression in female or male islets (**Figure 8A,G**), or on *Il1b* in male islets (**Figure 8G**). Interestingly, TCDD significantly increased *Il-1β* expression in βAhr^WT^ female islets but not βAhr^KO^ islets (**Figure 8A**), suggesting that TCDD-induced AhR activation in β-cells promotes inflammation in female islets. We did not observe any changes in % live (calcein+) (**Figure 8B,D,H,J**) or % dead (PI+) (**Figure 8C,D,I,J**) cells in dispersed islets of either sex. We also assessed β-cell apoptosis by immunofluorescence staining in pancreas sections at 1-week post-injection and found no changes in % TUNEL^-ve^ INS^+ve^ cells per islet (**Figure 8E,K**) or % INS^+ve^ area per islet (**Figure 8F,L**) in TCDD-exposed βAhr^WT^ and βAhr^KO^ mice of either sex.

**Figure 8:**
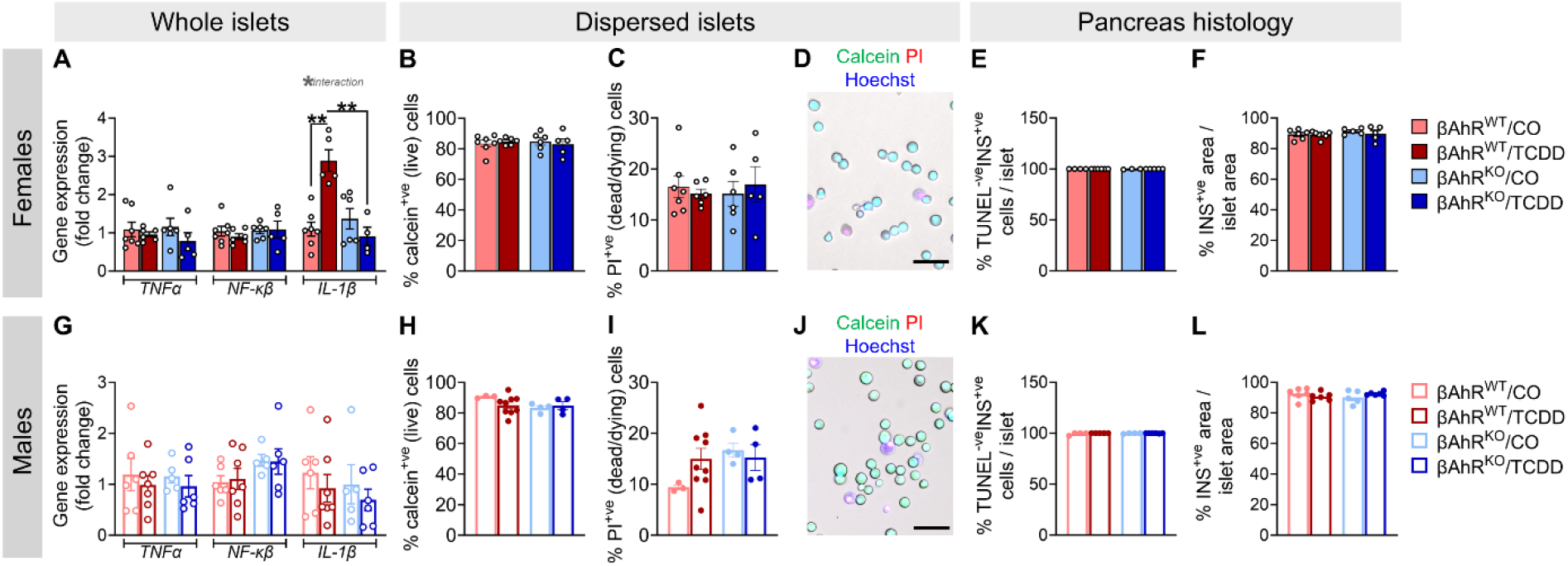
TCDD exposure increased *Il1b* expression in βAhr^WT^ but not βAhr^KO^ female islets. Islets were isolated at 1-week post-TCDD/CO injection to assess cell viability and gene expression, and whole pancreas was extracted for histological assessments (see Figure 3A for study timeline). **(A,G)** Gene expression for inflammatory markers in **(A)** female and **(G)** male islets (n = 5 – 7 mice / group). **(B,H)** % calcein^+ve^ (live) cells and **(C,I)** % propidium iodide (PI)^+ve^ (dead/dying) cells in islets from **(B,C)** female and **(H,I)** male mice (n = 3 – 9 mice / group; each mouse is an average of 1 – 2 technical replicates). **(D,J)** Representative images showing calcein, PI, and hoechst staining in dispersed islet cells. Scale bar = 50 μM. **(E,K)** % TUNEL^-ve^ insulin (INS)^+ve^ cells / islet, and **(F,L)** % INS^+ve^ area / islet area in pancreas sections from **(E,F)** female and **(K,L)** male mice (n = 4 – 7 mice / group). All data are presented as mean ± SEM. Individual data points in bar graphs represent biological replicates. *p ≤ 0.05, **p ≤ 0.01. The following statistical tests were used: two-way ANOVA with uncorrected Fisher’s LSD test.

We also measured markers of the AhR pathway in isolated islets at 1-week post CO/TCDD injection. TCDD caused an overall decrease in *Cyp1a1* in both sexes, irrespective of genotype (**Supplemental Figure 1A, B**). TCDD also caused a modest decrease in *Ahr* expression in βAhr^KO^ male islets compared to CO-exposed βAhr^KO^ islets (**Supplemental Figure 1B**), indicating negative feedback regulation of the AhR pathway in islets by 1 week post-injection. In line with our model validation data (**Figure 1F**), *Ahr* expression was downregulate in CO-exposed βAhr^KO^ female islets compared to βAhr^WT^ (**Supplemental Figure 1A**).

Lastly, we assessed whether high-dose TCDD exposure impacted gene expression of islet hormones and markers of β-cell function in βAhr^WT^ and βAhr^KO^ mouse islets at 6 weeks post CO/TCDD injection (**Supplemental Figure 2**). Despite the pronounced effects of TCDD on glucose homeostasis in βAhr^WT^ females, TCDD did not impact *Ins1*, *Ins2*, *Gcg*, *Pcks1*, *Psck2*, *Slc2a2*, *Gck*, or *MafA* expression in female islets, irrespective of genotype (**Supplemental Figure 3A,B**). In contrast, TCDD-exposure increased *Ins1, Gcg, and Slc2a2* expression in βAhr^WT^ and βAhr^KO^ male islets, but increased *Pcsk2* and *Gck* in βAhr^KO^ male islets only (**Supplemental Figure 2C,D**).

## 4. Discussion

Our study demonstrates that AhR signaling in β-cells plays a role in maintaining metabolic health in mice, even in the absence of exogenous chemical stimuli. βAhr^KO^ females had modestly reduced fasted blood glucose levels and increased basal and glucose-stimulated insulin secretion *ex vivo*, whereas βAhr^KO^ males displayed increase body weight compared to βAhr^WT^ controls. Overactivation of AhR in β-cells following chemical exposure appears to be detrimental for β-cell health. In line with our previous findings [24], we show that high-dose TCDD exposure causes fasting hypoinsulinemia in both female and male mice, but effects on glucose homeostasis are sex-specific. TCDD-exposed βAhr^WT^ females had reduced glucose-stimulated plasma insulin levels, insulin resistance, and pronounced hyperglycemia during a GTT; these effects were abolished in TCDD-exposed βAhr^KO^ females. TCDD-exposed βAhr^WT^ males showed reduced glucose-stimulated insulin secretion *in vivo* and *ex vivo*, increased insulin sensitivity, and hypoglycemia during a GTT, which was also mitigated by deleting Ahr in β-cells. Importantly, these data imply that AhR signaling in β-cells is not only driving the effects of TCDD on insulin secretion but also on peripheral insulin sensitivity in both sexes. Collectively, these data suggest that the loss of AhR (i.e. under baseline condition) or overstimulation of AhR (i.e. following chemical exposure) in β-cells impairs metabolic health, indicating the importance of this pathway in regulating β-cell physiology. Our study also points to a sex-specific role of AhR in β-cells.

We show convincing evidence that the detrimental effects of TCDD on glucose homeostasis and β-cell function are largely mediated by AhR signaling in β-cells. TCDD-exposed βAhr^WT^ female mice displayed insulin resistance and hyperglycemia, which was abolished by knocking out Ahr in β-cells. Interestingly, TCDD also increased *Il1b* expression in islets from βAhr^WT^ females but not βAhr^KO^ females at 1-week post-injection. It is well established that islet IL-1β signaling leads to β-cell dysfunction and β-cell apoptosis [29–32]. We did not observe an increase in islet cell apoptosis or changes in % insulin^+ve^ area per islet in female βAhr^WT^ mice at 1-week post-TCDD; however, it is plausible that persistent TCDD exposure (and thus, prolonged *Il1β* induction) would lead to β-cell apoptosis. Regardless, knocking out Ahr in female mice prevented the TCDD-induced increase in *Il1β* in islets and onset of glucose intolerance. These data suggest that AhR-mediated inflammation is likely contributing to β-cell dysfunction in female mice. However, TCDD-exposed females had reduced fasting plasma insulin levels irrespective of genotype, suggesting that AhR-independent mechanisms are mediating some of the effects of TCDD on glucose homeostasis.

In line with previous data [24], TCDD-exposed βAhr^WT^ males displayed reduced plasma insulin levels, increased insulin sensitivity, and hypoglycemia at 1-week post-injection, which coincided with reduced GSIS *ex vivo*. Past studies have speculated that TCDD alters glucose homeostasis primarily by increasing insulin sensitivity [33–35], which then causes adaptive changes in insulin secretion. Here we show that deleting Ahr in β-cells abolished all TCDD-induced metabolic phenotypes in males, including changes in peripheral insulin sensitivity, implying that β-cell dysfunction is likely driving metabolic phenotypes in TCDD-exposed male mice. In line with previous studies, TCDD-exposed βAhr^WT^ males also had reduced expression of insulin-regulated genes in liver [36–38]; we are the first to provide evidence that these well-documented effects of TCDD in liver are likely driven by changes in insulin secretion (given the absence of these changes in βAhr^KO^ male liver). The mechanism through which AhR activation impairs β-cell function in males remains unclear; we did not observe any changes in β-cell viability/apoptosis or inflammation genes in male islets. Previous studies have suggested that TCDD exposure can impair β-cell function by altering intracellular ATP levels [39] and/or Ca^2+^ influx [39,40]. Whether these effects are mediated by AhR signaling in β-cells requires further investigation.

We also provide novel evidence that AhR signaling in β-cells plays a role in maintaining metabolic health in the absence of chemical stimuli. βAhr^KO^ females had modestly reduced fasted blood glucose levels and increased basal and glucose-stimulated insulin secretion *ex vivo* compared to βAhr^WT^ females, suggesting that β-cell AhR signaling mediates β-cell function in female mice. However, it is important to note that βAhr^KO^ females displayed normal fasted and glucose-stimulated plasma insulin levels *in vivo*, potentially pointing to increased insulin clearance in βAhr^KO^ females; more detailed analyses are required to understand this phenotype. Interestingly, previous studies report normal fasting blood glucose, fasting c-peptide levels, and glucose tolerance in young global Ahr^KO^ female mice (3 – 4 months of age), but glucose intolerance, hypoinsulinemia, and insulin resistance in aged Ahr null female mice (7 – 15 months of age) [41,42]. Given that the subset of female mice that were maintained until 6 week post-injection in our study were ∼3-5 months of age, longer-term studies in βAhr^KO^ mice are warranted to better understand the role of AhR in β-cells. We also show that βAhr^KO^ males displayed increase body weight and reduced hepatic *Ppargc1a* expression compared to βAhr^WT^ controls. PGC1α is a master regulator of mitochondrial biogenesis, fatty acid oxidation, and energy substrate uptake/utilization [43]; PGC1α expression in liver has been negatively correlated with body fat content in humans [43] and liver-specific PGC1α knockout mice display weight gain [44]. As such, our data suggests a baseline role of β-cell AhR signaling in lipid homeostasis and/or body weight regulation. Future studies should characterise changes in fatty acid oxidation and lipid deposition in tissues such as adipose, as well as energy expenditure in βAhr^KO^ mice. It would also be interesting to investigate the role of β-cell AhR signaling in mediating energy homeostasis following HFD feeding.

Our findings in βAhr^KO^ mice under baseline condition largely differ from global βAhr^KO^ models. A study by Xu *et al*. [22] showed that when fed a chow diet, male Ahr null mice had normal body weight compared to WT mice but were protected from HFD-induced weight gain and hyperinsulinemia. In addition, Wang *et* al. [21] reported that global Ahr^KO^ mice had improved glucose tolerance and increased insulin sensitivity, but no change in glucose-stimulated plasma insulin levels; note male and female data were pooled in this study. Collectively, studies in global Ahr^KO^ mice suggest that loss of Ahr is protective against metabolic impairments, whereas our study suggests that loss of AhR in β-cells in the absence of a chemical stressor promotes weight gain and hypersecretion of insulin, which could be detrimental. These discrepancies between our data and studies in global AhR^KO^ null mice imply that loss of AhR signaling in β-cells may cause maladaptive AhR-dependent responses in peripheral tissues that are absent in global Ahr knockouts.

Our study provides novel evidence that AhR signaling in β-cells is required for maintaining β-cell function in female mice and body weight in male mice under baseline conditions. Additionally, we show that high-dose TCDD exposure impairs glucose homeostasis and β-cell function via AhR activation in β-cells. This implies that exposure to environmental AhR ligands directly targets the β-cell and can increase diabetes susceptibility, emphasizing the importance of considering the β-cell in chemical toxicity assessment.

## Abbreviations

AhR: Aryl hydrocarbon receptor
CO: Corn oil
Cyp1a1: Cytochrome P450 1A1
GTT: Glucose tolerance test
GSIS: Glucose-stimulated insulin secretion
Gsta1: Glutathione S-transferase alpha 1
Il-1β: Interleukin-1β
ITT: Insulin tolerance test
Nqo1: NAD(P)H: quinone oxidoreductase
POPs: Persistent Organic Pollutants
Ppargc1a: Peroxisome proliferative activated receptor 1α
TCDD: 2,3,7,8-tetrachlorodibenzo-p-dioxin
T2D: Type 2 diabetes

## Acknowledgements

This research was supported by a Canadian Institutes of Health Research (CIHR) Project Grant (#PJT-2018-159590), the Canadian Foundation for Innovation John R. Evans Leaders Fund (#37231), and an Ontario Research Fund award. M.P.H. was supported by a CIHR CGS-D award. M.E.A.C. was supported by an NSERC CGS-M and NSERC CGS-D award. L.B. was supported by an CIRTN-R2FIC-CREATE and CIHR CGS-D award. J.E.B. is supported by an Early Researcher Award from the Ontario Government.

## 6. Author Contributions

M.P.H. and J.E.B. conceived the experimental design, analysed the data, and wrote the manuscript. M.P.H., M.E.A.C., L.B., K.V.A, J.P., I.P., A.A.H., and J.E.B. were involved with data acquisition and interpretation of data. All authors contributed to manuscript revisions and approved the final version of the article.

