## Supplemental material for "The aryl hydrocarbon receptor in β-cells mediates the effects of TCDD on glucose homeostasis in mice"

**Supplemental Table 1:** Genotyping primer sequences.

| Target | Sequence (5'-3') | Amplicon Size (bp) |
|---|---|---|
| loxP-AhR Forward (#oIMR6075) | GGT ACA AGT GCA CAT GCC TGC | Wild-type = 106<br>Mutant = 140<br>Heterozygous = 106 and 140 |
| loxP-AhR Reverse (#oIMR6076) | CAG TGG GAA TAA GGC AAG AGT GA | |
| Cre Reverse (#26992) | GGA AGC AGA ATT CCA GAT ACT TG | Wild-type = 488<br>Mutant = 675<br>Heterozygous = 488 and 675 |
| Cre wild-type forward (#26993) | GTC AAA CAG CAT CTT TGT GGT C | |
| Cre mutant forward (#26994) | GCT GGA AGA TGG CGA TTA GC | |

**Supplemental Table 2:** *AhR* primers used to assess recombination.

| Target | Forward Sequence (5'-3') | Reverse Sequence (5'-3') | Amplicon Size (bp) |
|---|---|---|---|
| AhR Exon 1/3 primers | AAC ATC ACC TAT GCC AGC CG | CTG TCA GCA GGG GTG GAC TT | 263 |
| AhR Exon 2 primers | GCC CTT CCC GCA AGA TGT TA | AAG CTC TTG GCC CTC AGG TA | 84 |

**Supplemental Table 3:** Primer sequences for qPCR.

| Target | Forward Sequence (5'-3') | Reverse Sequence (5'-3') |
|---|---|---|
| *Acaca* | CCT GAC AAA CGA GTC TGG CT | CAT TCC ATG CAG TGG TCC CT |
| *Cyp1a1* | ATC ACA GAC AGC CTC ATT GAG C | AGA TAG CAG TTG TGA CTG TGT C |
| *G6pc* | TAC TAC AGC AAC AGC TCC GTG | TCC CAA CCA CAA GAT GAC GTT |
| *Gcg* | ACT CAC AGG GCA CAT TCA CC | CCA GTT TAT AAA GTC CCT GG |
| *Gck* | TGG TGG ATG AGA GCT CAG TG | TGA GCA GCA CAA GTC GTA CC |
| *Gsta1* | GCA AGG AAG GCT TTC AAG ATT CA | TTG CAA AAT AGC CAG GAT CAA CA |
| *IL-1β* | GCC ACC TTT TGA CAG TGA TGA G | AGC TTC TCC ACA GCC ACA AT |
| *Ins1* | TCA GAG ACC ATC AGC AAG CA | CTC CCA GAG GGC AAG CAG |
| *Ins2* | GCT TCT TCT ACA CAC CCA TGT | ACG ACT GAT CTA CAA TGC CAC |
| *MafA* | AGT CGT GCC GCT TCA AG | CGC CAA CTT CTC GTA TTT CTC C |
| *NF-Κβ* | CTC AGG AGC AGA AGT CTG GG | GCC GCT ATA TGC AGA GGT GT |
| *Nqo1* | CTC TGG CCG ATT CAG AGT GG | GTC TCC TCC CAG ACG GTT TC |
| *Pck1* | TGG GAA CTC ACT ACT CGG GA | TTC TTC TTG CCT TCG GGG TT |
| *Pcsk1* | GGT GGA AAG GTC GAG TCT AGC | TGC ACA CCA AAC GCA AAA GA |
| *Pcsk2* | TTT GGA GTC CGA AAG CTC CC | GGT GTA GGC TGC GTC TTC TT |
| *Ppargc1a* | CTT GAC TGG CGT CAT TCG G | CGC TAC ACC ACT TCA ATC CAC |
| *PPIA* | GCC AGG ACC TGT ATG CTT TA | AGC TCT GAG CAC TGG AGA GA |
| *Slc2a2* | GCA ACT GGG TCT GCA ATT TT | CCA GCG AAG AGG AAG AAC AC |
| *TNFα* | AGT CCG GGC AGG TCT ACT TT | ATG AAC ACC CAT TCC CTT CA |

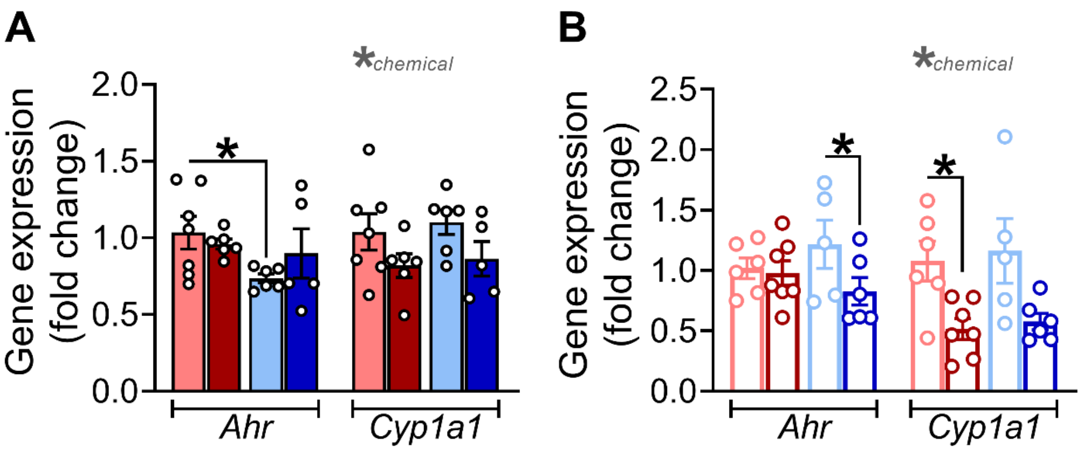


**Supplemental Figure 1: The AhR pathway is negatively regulated in islets by 1-week post-injection.** Islets were isolated at 1-week post TCDD/CO injection to assess expression of AhR pathway markers in **(A)** female and **(B)** male islets (n= 5 – 7 mice / group; see **Figure 3A** for study timeline). All data are presented as mean ± SEM. Individual data points in bar graphs represent biological replicates. *p ≤ 0.05, **p ≤ 0.01. The following statistical tests were used: two-way ANOVA with uncorrected Fisher's LSD test.

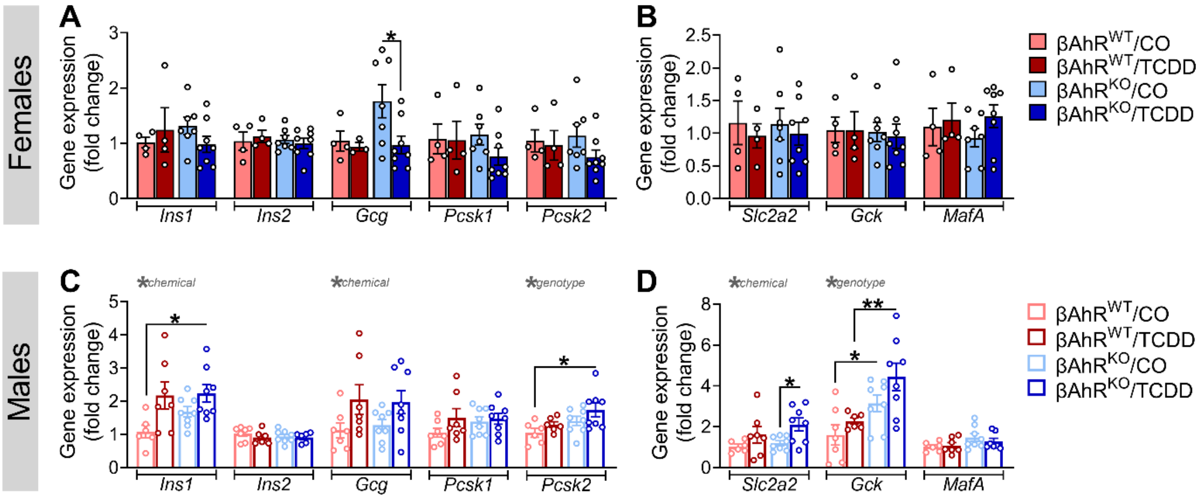


**Supplemental Figure 2: Knocking out Ahr in β-cells had modest effects on islet transcriptional changes in TCDD-exposed male mice.** Islets were harvested at 6 week post-TCDD/CO injection to assess gene expression of **(A,C)** hormones and hormone processing genes, and **(B,D)** markers of β-cell function in **(A,B)** female and **(C,D)** male mouse islets. All data are presented as mean ± SEM. Individual data points in bar graphs represent biological replicates (n = 4 – 8 mice / group). $^*p \le 0.05$, $^{**}p \le 0.01$. The following statistical tests were used: two-way ANOVA with uncorrected Fisher's LSD test.
